# Ankyrin G membrane partners drive the establishment and maintenance of the axon initial segment

**DOI:** 10.1101/073163

**Authors:** Christophe Leterrier, Nadine Clerc, Fanny Rueda-Boroni, Audrey Montersino, Bénédicte Dargent, Francis Castets

## Abstract

The axon initial segment (AIS) is a specialized neuronal compartment that plays a key role in neuronal development and excitability. It concentrates multiple ion channels and cell adhesion molecules. The anchoring of these AIS membrane components is known to be highly dependent of the scaffold protein ankyrin G (ankG) but whether ankG membrane partners play a reciprocal role in ankG targeting and stabilization has not been studied yet. In cultured hippocampal neurons and cortical organotypic slices, we found that shRNA-mediated knockdown of ankG membrane partners led to a decrease of ankG concentration and perturbed the AIS formation and maintenance. These perturbations were rescued by expressing an AIS-targeted sodium channel, or a minimal construct containing the ankyrin-binding domain of Nav1.2 and a membrane anchor. We thus demonstrate that a tight and precocious association of ankG to its membrane partners is crucial for the establishment and maintenance of the AIS.

## Introduction

Neurons exhibit an axonal/dendritic polarity that allows the directional propagation of signals throughout the nervous system. A major player in the maintenance of this polarity is the axon initial segment (AIS), a specialized domain located within the first 50 μm of the axon. The AIS isolates the axon from the cell body and regulates protein exchange between the axonal and somatodendritic compartments (Rasband, 2010; Leterrier, 2016). Besides this important role in neuronal polarity, the AIS also controls neuronal intrinsic excitability by generating action potentials (Clark et al, 2009; Kole & Stuart, 2012). At the molecular level, the AIS is organized by ankyrin G (ankG), a specialized scaffolding protein that directly binds to the submembrane cytoskeletal lattice of ßIV spectrin and actin. The aminoterminus of ankG, called the membrane-binding domain (MBD), is apposed to the inner face of plasma membrane and directly interacts with transmembrane proteins such as voltage-gated sodium channels (Nav), potassium channels Kv7.2/3 (KCNQ2/KCNQ3), and the cell adhesion molecules (CAMs) NrCAM and Neurofascin-186 (Nfasc186) (Leterrier et al, 2015). AnkG is also associated with the microtubule cytoskeleton by a direct interaction with End-Binding proteins (EB1 and EB3) (Leterrier et al, 2011b; Freal et al, 2016).

AnkG is considered to be the master organizer of the AIS (Rasband, 2010). It is the earliest component addressed to the AIS and is responsible for the subsequent recruitment of most AIS-enriched proteins (Jenkins, 2001; Hedstrom et al, 2008; Galiano et al, 2012). AnkG depletion not only impairs the accumulation of other AIS components, but also causes a progressive loss of neuronal polarity (Hedstrom et al, 2007; Sobotzik et al, 2009). Precisely how and where ankG interacts with its membrane partners during AIS formation and maintenance is still elusive. The “diffusion and trapping” model proposes that an existing ankG scaffold immobilizes membrane proteins at the AIS (Brachet et al, 2010; Xu & Cooper, 2015). Alternatively, recent data suggest that a preformed complex of ankG and Nav is transported to the AIS (Barry et al, 2014), and that Nav can be directly inserted at the AIS (Akin et al, 2015). Moreover, several results suggest an interplay between ankG and its membrane partners: the establishment of the AIS is impaired in motor neurons depleted for Nav (Xu & Shrager, 2005), and Nfasc186 elimination in Purkinje cells of adult mice results in progressive AIS disassembly (Zonta et al, 2011). In addition, we have shown that perturbing the ankG/Nav interaction by inhibiting of the protein kinase CK2 downregulates Nav accumulation at the AIS, and subsequently decreases ankG concentration (Brechet et al, 2008; Brachet et al, 2010). To reveal and characterize the interplay between ankG and its membrane partners, here we have examined the role of AIS membrane proteins (Nav and Nfasc186) for AIS assembly and maintenance. Knockdown of Nav and Nfasc-186 resulted in a cumulative impairment of the AIS establishment. Rescue with a full-length Nav1.6 or a minimal construct combining the ankG binding domain of Nav and a membrane targeting motif showed that anchoring of ankG to the plasma membrane via its partners is necessary for its targeting and stable assembly at the AIS. Finally, overexpression of the membrane-anchored ankG-binding construct induced a mislocalization of ankG, suggesting that interaction of ankG with its membrane partners occurs before the insertion of ankG into the AIS.

## Results

### Nav1 knockdown reduces Na+ current in organotypic slices and impairs ankG concentration

We and others have demonstrated that Nav1 clustering at the AIS requires a direct interaction between the ankyrin-binding-domain (ABD) of Nav channels and the MBD of ankG (Garrido et al, 2003; Gasser et al, 2012; Montersino et al, 2014). We wanted to know if the Nav-ankG interaction could conversely contribute to ankG concentration and AIS integrity. We first assessed the effect of Nav depletion on ankG concentration at the AIS in cultured organotypic slices (Stoppini et al, 1991). Cortical slices obtained from 7 day rats were transduced with a recombinant lentivirus expressing a previously validated shRNA directed against Nav1.1/Nav1.2/Nav1.3 (shNav; Xu & Shrager, 2005; Hedstrom et al, 2007; Hien et al, 2014) or against ankG (shAnkG; Hedstrom et al, 2007; Leterrier et al, 2011). After 7 days, we evaluated the AIS integrity using immunostaining against Nav and ankG. The membrane-targeted GFP (m GFP) marker co-expressed with the shRNA allowed visualizing the full morphology of the transduced neurons (Figure 1). AnkG staining was readily observable in 86% of the neurons transduced with a control shRNA (Figure 1A & 1D) but was detected in only a small proportion of neurons expressing shAnkG (Figure 1B & 1D). In neurons transduced with shNav, ankG accumulation was observed in only 48% of the neurons (Figure 1C & 1D). As compared to control neurons, Nav1-depleted neurons had a significantly reduced ankG concentration at the AIS (ankG ratio - mean intensity of ankG labeling at the AIS and normalized to ankG labeling in surrounding non-infected neurons - down from 1.10±0.06 in shCtrl neurons to 0.72±0.05 in shNav neurons, Figure 1E). These results demonstrate that the depletion of Nav induces a significant decrease of ankG accumulation at the AIS.

To evaluate how Na+ currents were affected by ankG or Nav silencing, we performed electrophysiological recording on cultured slices, 8 to 9 days after infection. In large neurons with fully developed processes, space clamp problems prevent adequate voltage clamp recording of rapidly inactivating, i.e. transient, Na+ currents (INaT). However, all the Nav types expressed in central nervous system neurons generate also a slowly inactivating, i.e. persistent, Na+ current (INaP) that follows INaT (Mantegazza et al, 2005; Rush et al, 2005; Estacion & Waxman, 2013). Using depolarizing voltage ramps at adequate speed, we were able to record both correctly clamped INaP and unclamped INaT (Del Negro et al, 2002). The recorded currents were suppressed in the presence of TTX, showing that TTX-sensitive Nav channels generate these currents. In all the neurons transduced with the control shRNA, voltage ramps generated a large INaP that started to activate at −58.2 ± 1.8 mV (Figure 2E) and exhibited usually 2 components (Figure 2A), with the largest peaking at −33.9 ± 2.7 mV (Figure 2F). As expected, in most of these neurons (13/19), unclamped INaT were also generated and appeared as a single spike (star in Figure 2A; average threshold: −48.5 ±1.4 mV; average peak amplitude: 2467 ± 648 pA). In neurons expressing shAnkG, INaP was affected both in terms of amplitude (strong reduction in 4/5 neurons and null current in 1/5 neuron; Figure 2B & 2D) and in terms of voltage dependence (depolarizing shift of the activation threshold and peak voltage; Figure 2E & F). In addition, in these ankG depleted neurons, INaT was in most cases absent (4/5 neurons) or extremely small (1/5 neurons, trace presented in Figure 2B; threshold: −39 mV; peak amplitude: 208 pA). In the neurons expressing shRNA against Nav, INaP exhibited significant changes in voltage dependence (more depolarized threshold and peak voltage, Figure 2E & 2F), but its average amplitude was not significantly lower than in shCtrl neurons (Figure 2D). Remarkably, INaT was absent in all the Nav-depleted neurons (9/9 neurons). Altogether these results confirm that Nav channels were functionally eliminated in our shRNA experiments.

**Figure 1.**
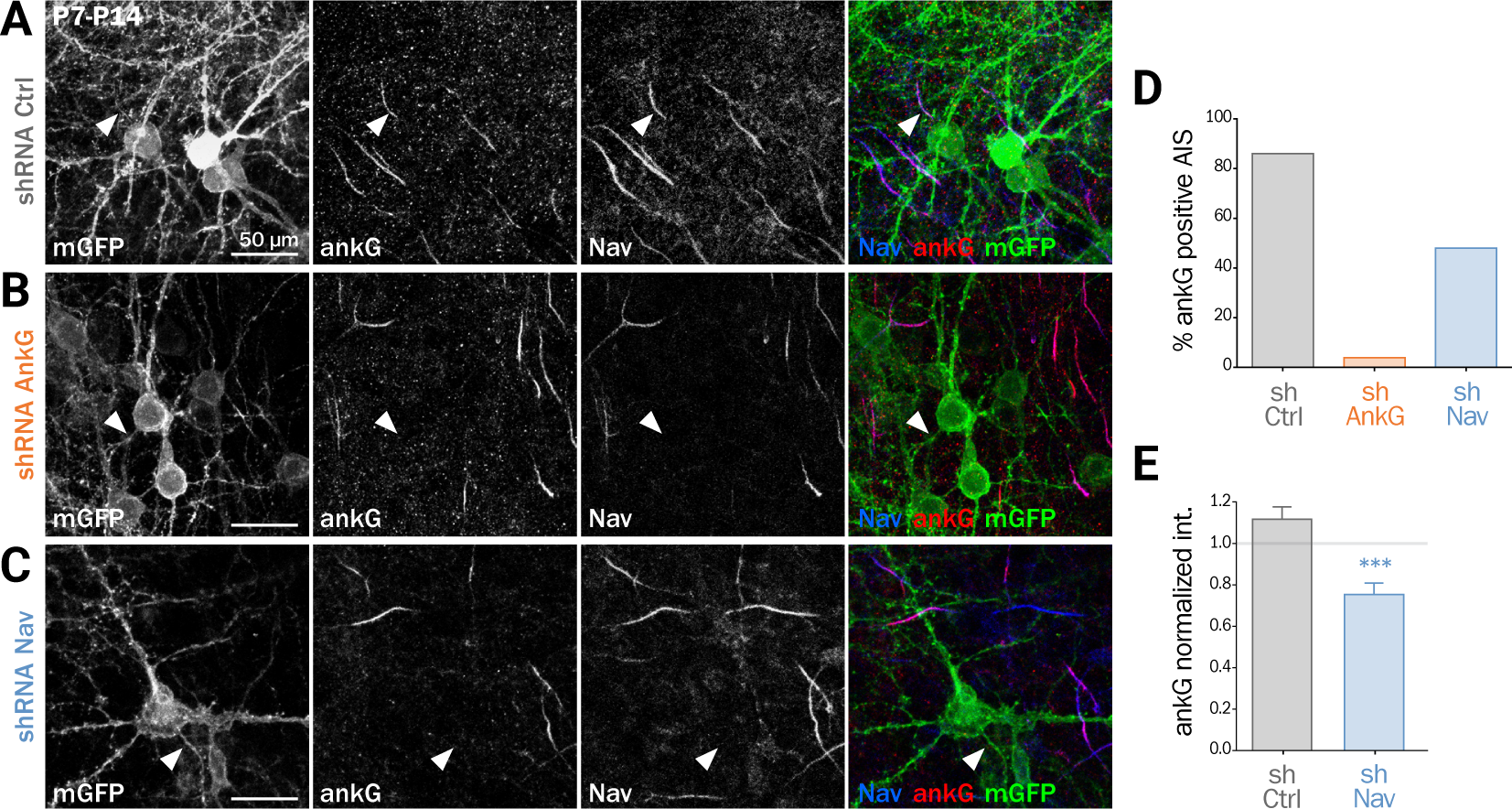
Knockdown of Nav channels downregulates the AIS in organotypic slices. Organotypic cortical slices prepared from a postnatal day 7 rat were infected with a lentivirus co-expressing membranous GFP (mGFP) and a shRNA directed against ankG (shAnkG), Nav (shNav) or luciferase as control (shCtrl). 7 days post-infection, slices were fixed, immunostained for GFP, ankG and Nav. **A, B and C:** Maximum intensity projection of 15−20 optical slices. Scale bar: 50ìm. D: Percentage of infected neuron with an observable AIS (shCtrl n=80; shAnkG n=27; shNav n=97). **E:** Ratio of the mean fluorescence intensity for ankG labeling at the AIS in transfected neurons (where the AIS was still detected) compared to the surrounding untransfected cells (shCtrl 1.10 ± 0.06, n=69; shNav: 0.72 ± 0.05, n=47; 3 independent experiments).

### Nav or Nfasc 186 knockdown impair AIS maintenance in cultured neurons

To decipher the mechanisms involved in the interplay between Nav and ankG, we tested the effect of Nav depletion in cultured hippocampal neurons, an amenable model for the analysis of neuronal morphology and polarity (Kaech & Banker, 2006). Neurons were maintained for 8 days in vitro (8 div), ensuring that polarity was well established, then transfected with shRNA constructs, and fixed 6 days later (14 div)(Figure 3). We first verified that the ankG labeling intensity was strongly reduced in shAnkG transfected neurons, compared to untransfected surrounding neurons (Figure 3B): ankG intensity ratio between transfected and untransfected neurons was 0.97±0.03 for shCtrl, but only 0.15±0.01 for shAnkG (Figure 3F). Similarly, we checked the efficiency of Nav depletion and found that the Nav intensity ratio was 0.96±0.04 for shCtrl and 0.37±0.04 for shNav (Figure 3E). As expected, AnkG depletion also resulted in Nav disappearance (Nav ratio 0.35+0.03, Figure 3E). Consistently with our results in slices, Navi depletion was accompanied by a ∼50% decrease in ankG labeling in the AIS (ankG ratio 0.48+0.04, Figure 3F). Thus, Navi expression is required for ankG concentration at the AIS, and therefore for AIS maintenance. We reasoned that Nav could have a stabilizing effect by anchoring ankG to the membrane. Since the MBD of ankG can bind both to Nav and CAMs such as Nfasc186, we examined whether Nfasc186 can also participates in ankG stabilization at the AIS, as suggested by data obtained after conditional genetic depletion (Zonta et al, 2011). Six days after transfection of 8 div neurons with a validated shRNA against Nfasc186 (Hedstrom et al, 2007), ankG accumulation at the AIS was quantitatively analyzed. AnkG labeling was significantly reduced in neurons depleted for Nfasc186 (1.02+0.05 for shCtrl, 0.59+0.03 for shNF, Figure 3D & 3G). This additional observation suggests that elimination of membrane partners of ankG (Nav or CAMs) is sufficient to induce AIS destabilization in polarized neurons.

**Figure 2.**
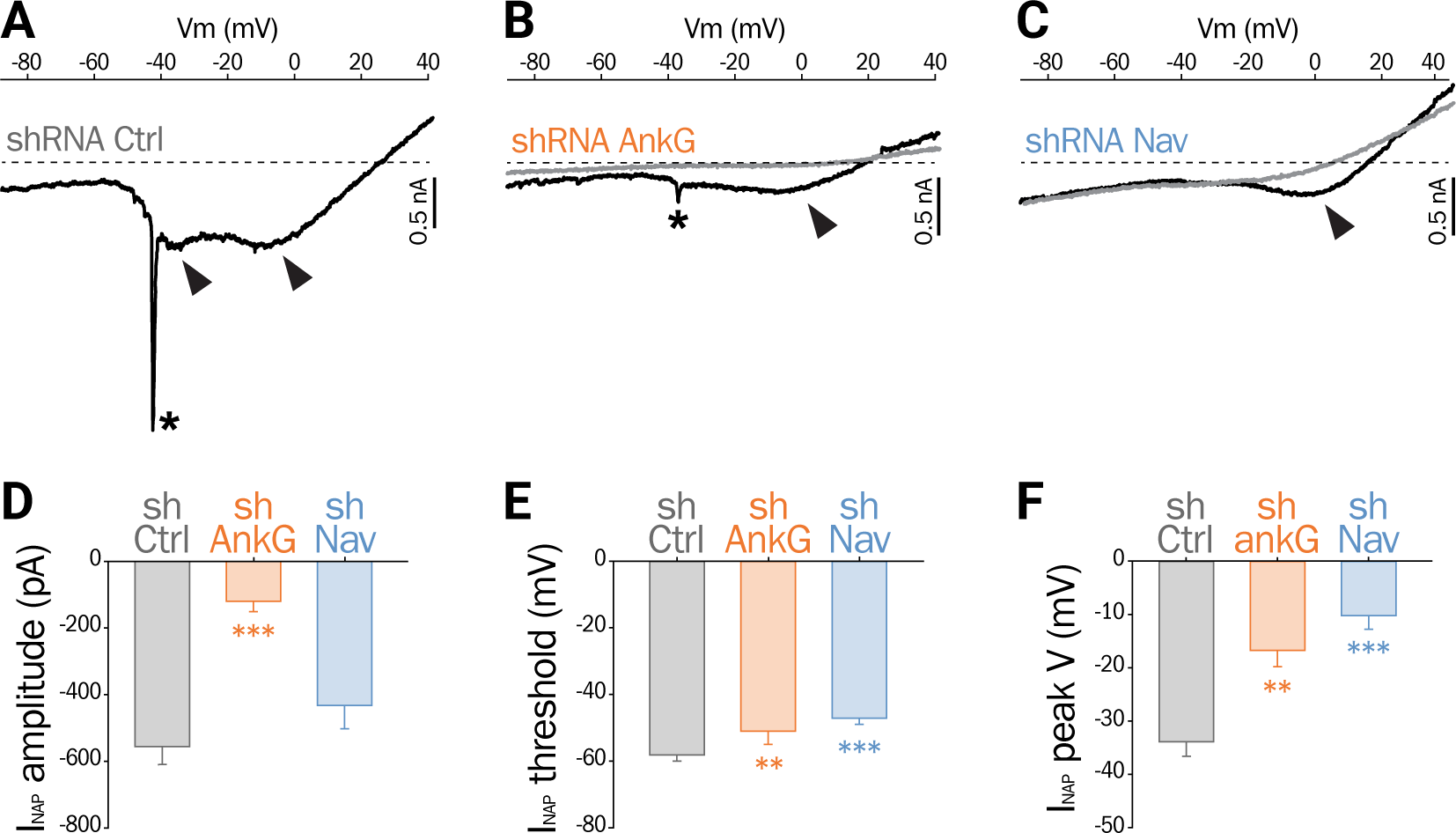
Effect of ankG and Nav knockdown on sodium currents. mGFP expressing neurons from organotypic cortical slices were analyzed by patch-clamp recording. **A, B and C**: Raw traces of Na+ currents evoked by voltage ramps (from holding potential −90 mV to +40 mV at 0,2 mV/ms) in the absence (black) or presence (grey) of 500 nM TTX. Asterisks and arrows respectively indicate transient Na+ currents and components of the Na+ persistent current (INaP). **D, E and F**: Characteristics of the INaP in the different conditions (shCtrl n=19; shAnkG n=5; shNav n=9). D: averaged peak amplitude; E: peak threshold; F: maximum peak voltage.

**Figure 3.**
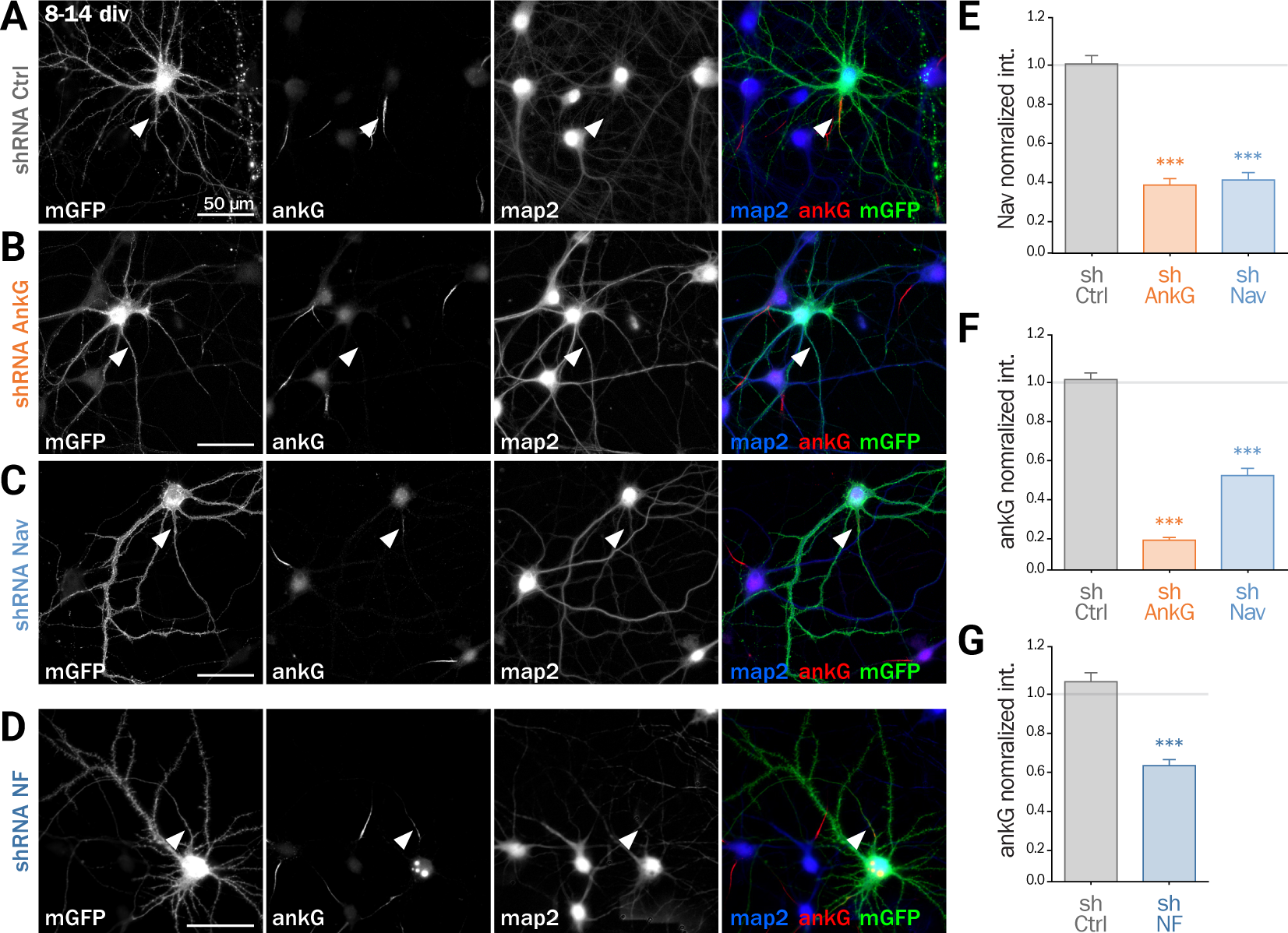
Knockdown of AIS membrane components perturbs AIS maintenance. **A, B, C and D:** Cultured rat hippocampal neurons were transfected at 8 div with mGFP and shCtrl (A), shAnkG (B), shNav (C) or shNF (D). 6 days later (14 div), neurons were fixed and immunostained for GFP, ankG, map2 and Nav. Arrowheads indicate the AIS of transfected neurons. E: Ratio of the mean fluorescence intensity for Nav1 labeling at the AIS in transfected neurons compared to surrounding untransfected cells (shCtrl 0.96 ± 0.04, n=43; shAnkG 0.35 ± 0.03, n=43; shNav 0.37 ± 0.04, n=40; 3 independent experiments). **F and G:** Ratio of the mean fluorescence intensity for ankG labeling at the AIS in transfected neurons compared to surrounding untransfected cells (F: shCtrl 0.97 ± 0.03, n=116; shAnkG n=0.15 ± 0.01, n=94; shNav 0.48 ± 0.04, n=123; 6 independent experiments. G: shCtrl 1.02 ± 0.05, n=36; shNF 0.59 ± 0.03, n=64; 3 independent experiments).

**Figure 4.**
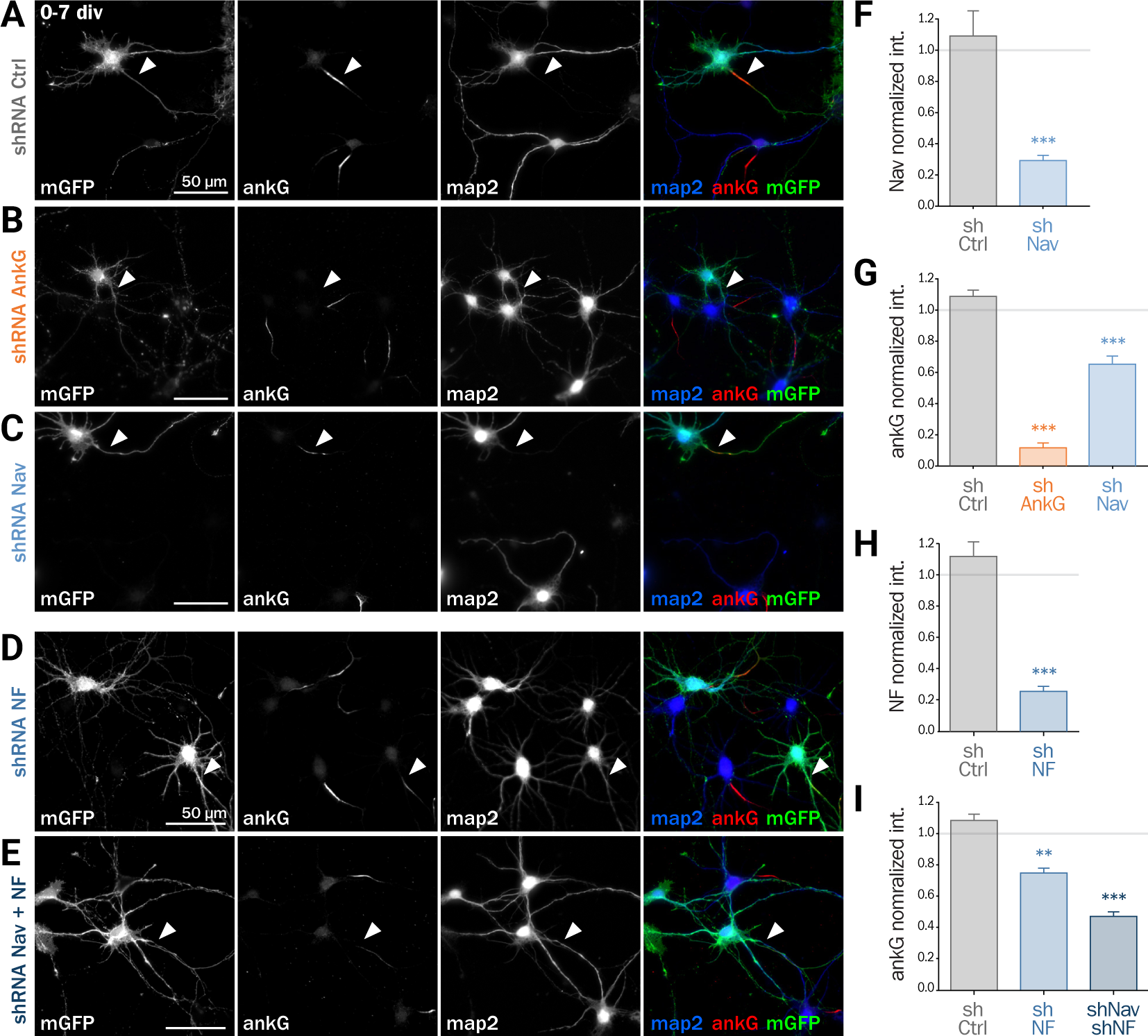
Knockdown of AIS membrane compononts impairs AIS formation. **A, B, C, D and E:** Rat hippocampal neurons were transfected before plating (0 div) with mGFP and shCtrl (A), shAnkG (B), shNav (C), shNF (D) or shNav+shNF (E). 7 days later (7 div), neurons were fixed and immunostained for GFP, ankG, map2 and Nav or Nfasc186. Arrowheads indicate the AIS of transfected neurons. **F:** Ratio of the mean fluorescence intensity for Nav1 labeling at the AIS in transfected neurons compared to surrounding untransfected cells (shCtrl 1.08 ± 0.12, n=41; shNav 0.30 ± 0.03, n=55; 3 independent experiments). **G:** intensity for ankG labeling at the AIS in transfected neurons compared to surrounding untransfected cells (shCtrl 1.08 ± 0.04, n=80; shAnkG 0.12 ± 0.03, n=40; shNav 0.65 ± 0.05, n=39; 3 independent experiments). **H:** Ratio of the mean fluorescence intensity for Nfasc186 labeling at the AIS in transfected neurons compared to surrounding untransfected cells (shCtrl 1.12 ± 0.09, n=20; shNF 0.25 ± 0.03, n=25; 2 independent experiments). **I:** Ratio of the mean fluorescence intensity for ankG labeling at the AIS in transfected neurons compared to surrounding untransfected cells (shCtrl 1.08 ± 0.04, n=80; shNF 0.75 ± 0.03, n=43 shNav+shNF 0.47 ± 0.03, n=109; 3 independent experiments).

### Nav or Nfasc 186 knockdown impairs AIS formation

Next, we assessed if ankG membrane partners are also required for proper AIS formation in young neurons. In developing hippocampal neurons in culture, polarity is established around 2 div and is quickly followed by AIS assembly (Dotti et al, 1988; Hedstrom et al, 2007). Freshly dissociated hippocampal neurons were nucleofected in order to express shRNA prior to AIS formation and then fixed after 7 div. We first checked that the expression of each shRNA led to the specific down regulation of its corresponding targets: ankG (ankG ratio 1.08±0.04 for shCtrl; 0.12±0.03 for shAnkG, Figure 4G), Nav1 (Nav ratio 1.08±0.12 for shCtrl; 0.30±0.03 for shNav, Figure 4F) or Nfasc186 (Nfasc186 ratio 1.12±0.09 for shCtrl, 0.26±0.03 for shNF, Figure 4H). As observed in mature neurons, ankG concentration was impaired in neurons depleted for either Nav channels (ankG ratio 0.65 ± 0.05 for shNav, Figure 4G) or Nfasc186 (ankG ratio 0.75 ± 0.03 for shNF, Figure 4I). In addition, we found that the effects of Nav or Nfasc186 depletion on AIS formation were cumulative, as simultaneous depletions using both shRNAs (shNav+shNF) further reduced ankG concentration at the AIS (0.47 ± 0.03, Figure 4I). Altogether, these data demonstrate that the membrane partners of ankG, Nav and Nfasc186, are essential for AIS assembly.

### Nav1.6-GFP expression rescues the ankG downregulation induced by depleting Nav or Nfasc186

If Nav specifically contribute to AIS formation and integrity, ankG downregulation induced by Nav depletion should be rescued by the co-expression of an shRNA-resistant Nav construct. To test this hypothesis, we used a full length Nav1.6-GFP that is resistant to our shRNA construct against Nav (3 bases mismatch)(Gasser et al, 2012). Nav1.6-GFP and a shNav plasmid containing soluble Td-Tomato as transfection marker were co-expressed either in freshly dissociated or mature (8 div) hippocampal neurons that were analyzed 6−7 days later. By contrast with mGFP that filled the entire neuron, Nav1.6-GFP was concentrated in the soma and at the AIS of co-transfected cells (Figure 5A-C). Quantitative analysis showed that ankG concentration was restored at the AIS by Nav1.6-GFP co-expression either in young neurons (0.56±0.07 for shNav+mGFP, 1.09±0.08 for shNav+Nav1.6-GFP, Figure 5F) or in mature neurons (0.37±0.05 for shNav+mGFP, 0.92±0.06 for shNav+Nav1.6-GFP, Figure 5G).

If ankG stabilization depends on ankG membrane anchoring per se and not on the identity of partners, overexpression of Nav1.6-GFP could also be able to cross-rescue Nf186 depletion (Figure 5D & 5E). When Nav1.6-GFP was co-expressed with the shRNA against Nfasc186, we indeed observed that ankG concentration at the AIS was restored in both young (0.68±0.03 for shNF+mGFP, 1.09±0.04 for shNF+Nav1.6-GFP, Figure 5H) and mature neurons (0.59±0.03 for shNF+mGFP, 0.95±0.04 for shNF+Nav1.6-GFP Figure 5I). This cross-rescue experiment demonstrates that one ankG membrane partner can be replaced by another and that the identity of the partner is less important that the membrane anchoring itself, to drive AIS formation and maintenance.

**Figure 5.**
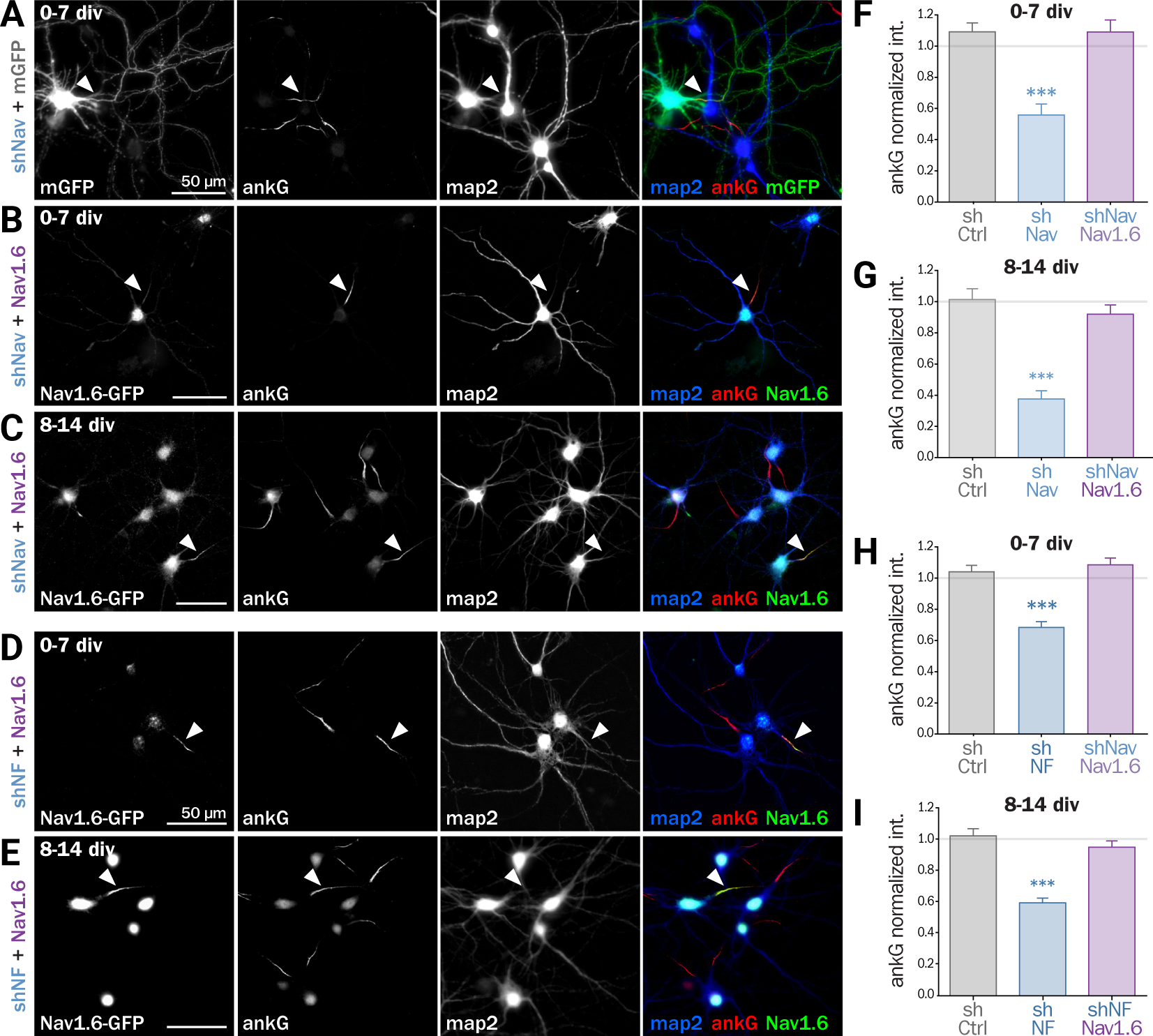
Nav1.6 rescue the AIS down-regulation induced by Nav knockdown, but also by Nfasc186 knockdown. **A, B C, D and E:** Rat hippocampal neurons were co-transfected at 0 div or 8 div with mGFP or Nav1.6-GFP and shNav (A, B and C) or shNF (D and E). 6 or 7 days later (7 div or 14 div), neurons were fixed and immunostained for GFP, ankG and map2. Arrowheads indicate the AIS of transfected neurons. **F, G, H and I:** Ratio of the mean fluorescence intensity for ankG labeling at the AIS in transfected neurons from 0 to 7 div (F and H) or 8 to 14 div (G and I) compared to surrounding untransfected cell. (F: shCtrl 1.09 ± 0.06, n=54; shNav 0.56 ± 0.07, n=21, shNav+Nav1.6 1.09 ± 0.07, n=41; 3 independent experiments. **G:** shCtrl 1.01 ± 0.07, n=28; shNav 0.37 ± 0.05, n=39; shNav+Nav1.6 0.92 ± 0.06, n=30; 2 independent experiments. H: shCtrl 1.04 ± 0.04, n=61; shNF 0.68 ± 0.04, n=64; shNF+Nav1.6 1.09 ± 0.04, n=68; 3 independent experiments. I: shCtrl 1.02 ± 0.05, n=36; shNF 0.59 ± 0.03, n=64; shNF+Nav1.6 0.95 ± 0.04, n=58; 3 independent experiments).

### Expression of a chimeric membrane-anchored ankG partner rescues AIS formation

Next, we wanted to directly prove that ankG association to the membrane via a protein partner contributes to ankG targeting and assembly. We devised a minimal chimeric protein bearing the well-described ABD from the intracellular loop II-III of Nav1.2 (Garrido et al, 2003; Lemaillet et al, 2003; Gasser et al, 2012) fused to the farnesylated form of GFP that associates to the plasma membrane (Figure 6A & 6B). When this membrane ABD (mABD) construct was expressed in young hippocampal neurons, it was highly concentrated along the proximal axon with an AIS/dendrite ratio of 5.46±0.53 (Figure 6C & 6F). As ABD binding to ankG is known to be impaired by the mutation of glutamate residue Nav1.2 E1111 or the four serine residues implicated in CK2 regulation (Brechet et al, 2008), we produced three mutant constructs where E1111, the four serine residues or all these five amino-acids were replaced by alanine (mABD-EA, mABD-4SA and mABD-E4S; Figure 6B). These mutations abolished the ability of mABD to be concentrated to the AIS, resulting in an AIS/dendrite ratio close to 1 (Figure 6F). This indicates that mABD concentration at the AIS is under the control of a phospho-dependent interaction with ankG. We next assessed whether the expression of mABD was able to rescue the ankG downregulation induced by either Nav or Nf186 depletion. In young neurons, mABD expression rescued the deficit in ankG accumulation caused by depletion of either Nav (ankG ratio 1.47 ± 0.15 for shNav+mABD; Figure 6D & 6GF) or Nfasc186 (ankG ratio 1.27 ± 0.07 for shNF+mABD, Figure 6E & 6H). Notably, we observed that over-expression of mABD by itself up-regulated AIS components such as ankG, β4-spectrin and Nfasc186 (ratios of 1.50±0.04 for ankG, 1.52±0.09 for β4 spectrin, and 1.62±0.14 for NF, Figure 6I, 6J & 6L). By opposition, endogenous Nav were strongly downregulated, indicating that the ABD domain acts as a dominant negative on endogenous Nav targeting or anchoring during AIS formation (Nav ratio 0.67±0.04, Figure 6K & 6L). In addition, mABD expression affected the morphology of the AISs that were wider, but not longer, than those of untransfected neurons (length 17.7±0.8 μm for untransfected, 18.4±0.7 μm for mABD, width 0.76±0.02 μm for untransfected, 1.78±0.07 pm for mABD, Figure 6M & 6N). Altogether, these experiments in young neurons during AIS formation show that mABD is able to rescue the absence of Nav or Nfasc186, and that its overexpression upregulates the whole AIS assembly, leading to a wider AIS that accumulate a higher density of components at the exception of Nav channels.

**Figure 6.**
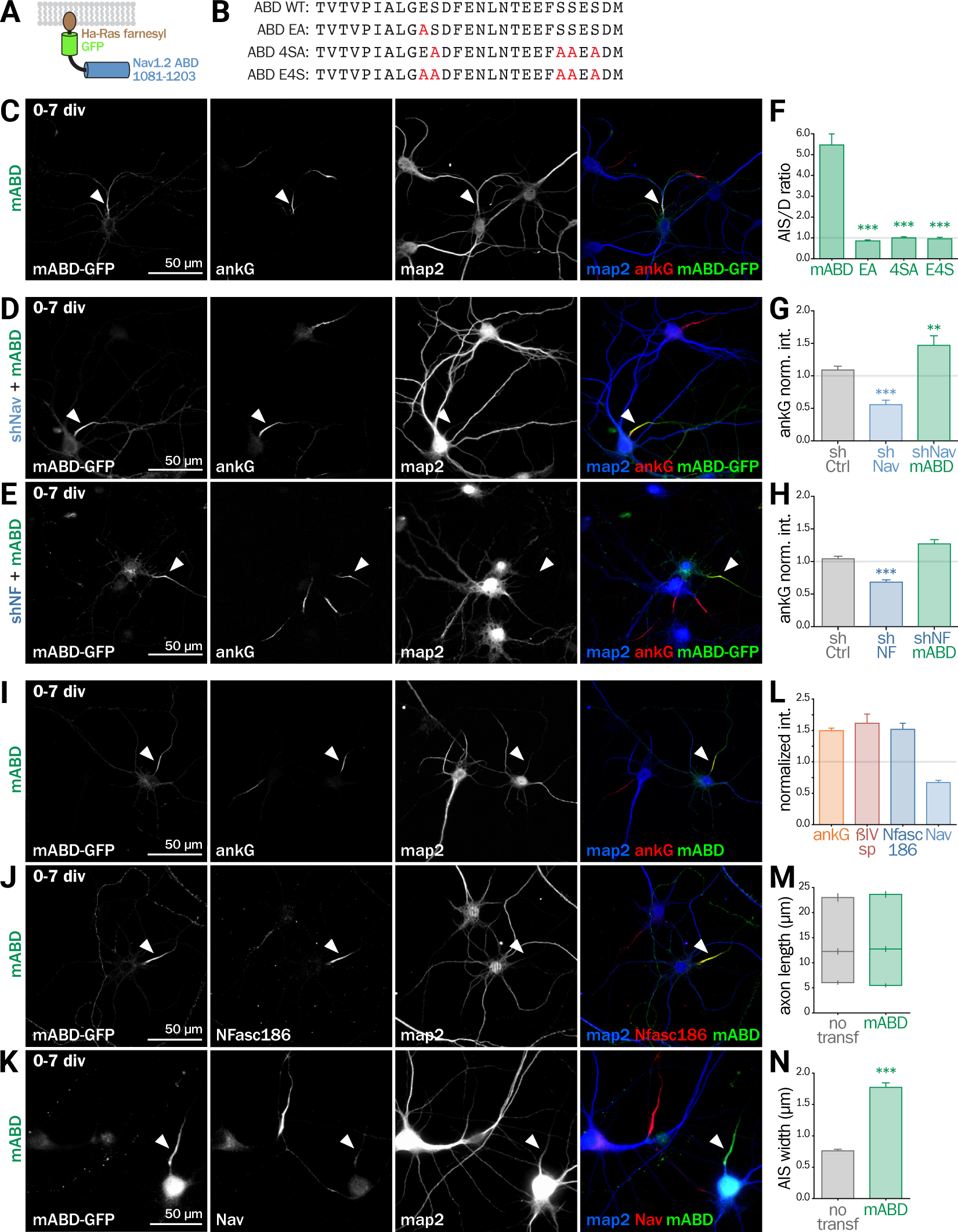
Synthetic mABD construct rescues Nav or Nfasc186 knockdown and upregulates AIS formation. **A:** Schematic representation of the mABD construct: amino-acids 1081-1203 of rat Nav1.2 were fused to GFP followed by a farnesylation motif from Ha-Ras. **B:** Alignment of wild type (WT) and mutated sequences corresponding to the rat Nav1.2 (aa 1102-1128) Ankyrin Binding Domain (ABD). The mutated amino-acids involved in ankG binding are in red. **C, D and E:** Rat hippocampal neurons were transfected at 0 div with mABD alone (C) or in association with shNav (D) or shNF (E). 7 days later (div 7), neurons were fixed and immunostained for GFP, map2 and ankG (C, D, E). Arrowheads indicate the AIS of transfected neurons. **F:** Ratio of the mean fluorescence intensity for GFP labeling in the proximal axon compared to the mean of three dendrites (D) of the same transfected neuron (mABD 5.46 ± 0.54, n=77, mABD-EA 0.86 ± 0.06, n=37, mABD-4SA 1.01 ± 0.05, n=36; mABD-E4S 0.96 ± 0.07, n=32; 2 independent experiments). **G and H:** Ratio of the mean fluorescence intensity for ankG labeling at the AIS in transfected neurons compared to surrounding untransfected cells (G: shCtrl 1.09 ± 0.06, n=54; shNav 0.56 ± 0.07, n=21; shNav+mABD 1.47 ± 0.15, n=23; 2 independent experiments. H: shCtrl 1.04 ± 0.04, n=61; shNF 0.68 ± 0.04; n=64; shNF+mABD 1.27 ± 0.07, n=49; 3 independent experiments). **I, J and K:** Rat hippocampal neurons were transfected at 0 div with mABD alone. 7 days later (7 div), neurons were fixed and immunostained for GFP, map2 and ankG (I), Nfasc186 (J), Nav (K), or ßIV-spectrin (not pictured). Arrowheads indicate the AIS of transfected neurons. **L:** Ratio of the mean fluorescence intensity for ankG, βIV-spectrin (βIVsp), Nfasc186 (NF) or Nav labeling at the AIS in transfected neurons compared to surroundings untransfected cells (ankG 1.50 ± 0.04, n=146; ßIVsp 1,52 ± 0.10, n=49; NF 1,62 ± 0.14, n=26; Nav0.67 ± 0.04; n=33; 2 to 6 independent experiments). **M and N:** Length and the width of the AIS in mABD-transfected neurons (untransfected cells AIS width 0.76 ± 0.03 μm, n=61; mABD 1.78 ± 0.07 μm, n=60; 2 independent experiments).

### mABD expression in mature neurons results in ankG ectopic localization

Next, we assessed the effect of mABD expression in mature neurons. First, we turned back to post-natal cortical organotypic slices, and examined neurons transduced with mABD (together with a control shRNA). In mature neurons with an already assembled AIS, mABD was localized in the whole neuron, with no concentration at the AIS (Figure 7A). This contrasted with the AIS localization observed when expressing mABD during AIS formation in cultured neurons (see above). Interestingly, this non-polarized expression of mABD was accompanied by a partial delocalization of ankG, which was found to accumulate in the soma and proximal dendrites in addition to the AIS (Figure 7A). This was evidenced by the drop of the AIS/soma polarity index for ankG labeling in neurons expressing mABD compared to neurons expressing the neutral marker mGFP (0.88±0.03 for mGFP, 0.14±0.09 for mABD, Figure 7C). We reasoned that this difference in mABD localization during AIS formation or maintenance could be explained by the presence, in the AIS of mature neurons, of strongly anchored endogenous Nav that compete with the mABD construct for ankG binding sites. To test this hypothesis, the mABD together with the shRNA against Nav channels were expressed in organotypic slices. In contrast to control neurons, in the Nav-depleted neurons, the mABD was concentrated to the AIS and did not delocalize ankG (Figure 7B & 7C)(ankG polarity index for mABD+shNav: 0.79 ±0.03). Thus, the expression of mABD in mature neurons depleted for Nav results in a proper stabilization of ankG at the AIS. This also means that mABD expression in organotypic slices rescues the ankG downregulation observed in neurons depleted for Nav (see Figure 1).

**Figure 7.**
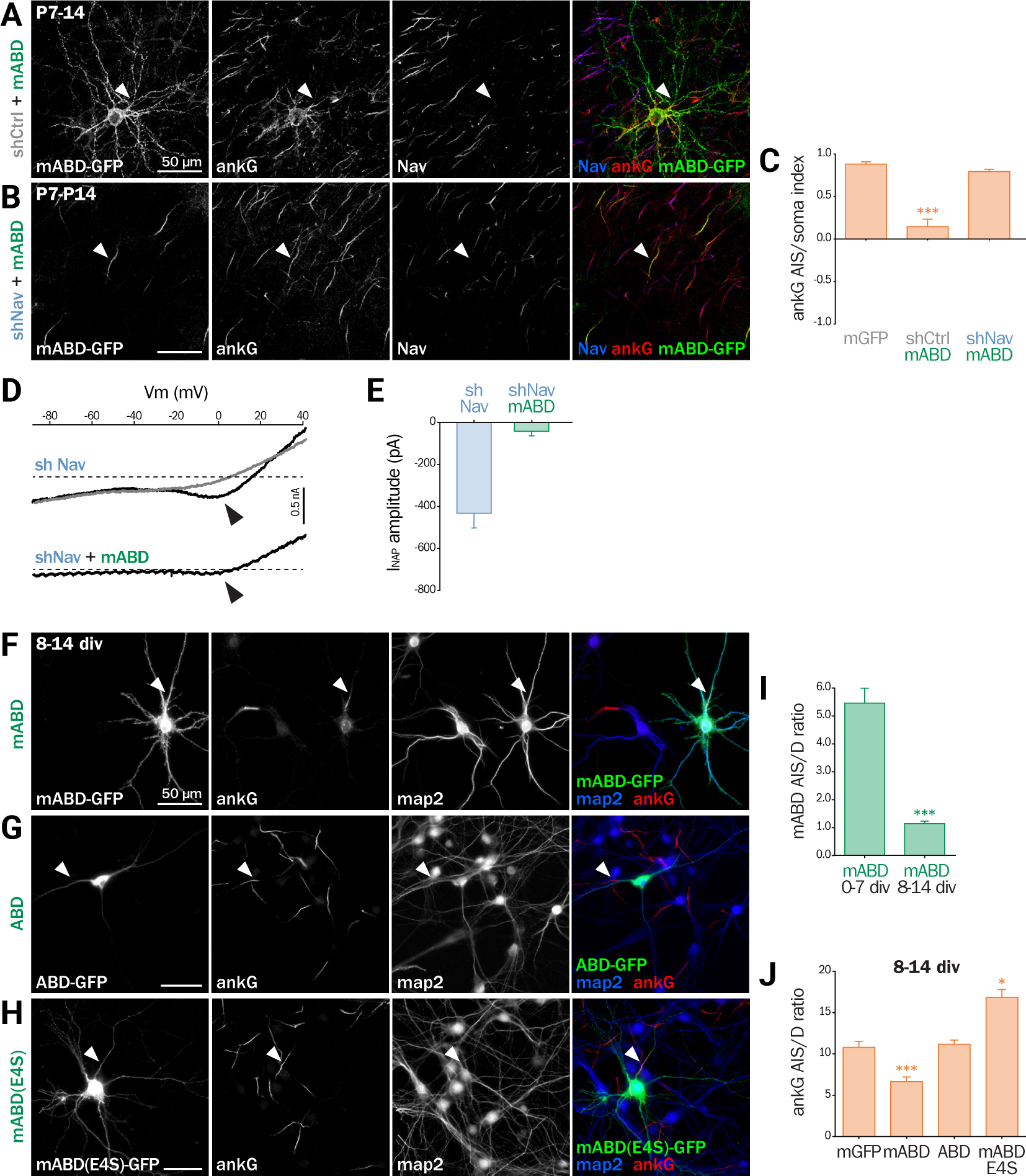
mABD expression in mature neurons results in ankG mislocalization. **A, B:** Organotypic cortical slices from postnatal day 7 rats were infected with a lentivirus co-expressing mABD and shCtrl or shNav. 7−8 days post-infection, slices were either fixed, immunostained for GFP, ankG and Nav. Maximum-intensity projection of 15−20 optical slices. Arrowheads indicate the AIS of transfected neurons. **C:** Polarity index of ankG labeling in the AIS versus the soma of infected neurons (mGFP alone 0.88 ± 0.03, n=8; mABD+ shCtrl 0.15 ± 0.09, n=10; mABD+shNav 0.79 ± 0.03, n=10; 2 independent experiments). **D and E:** Organotypic slices similar to A-C were analyzed by patch-clamp recording. D, Raw traces of Na+ currents evoked by voltage ramps in the absence (black) or presence (grey) of 500 nM TTX. Arrowheads indicate the Na+ persistent current (INaP). E, Averaged peak amplitude (shNav n=9; shNav+mABD: n=3/7). **F, G and H:** Rat hippocampal neurons were transfected at 8 div with mGFP, mABD, ABD (mABD devoid of its membrane anchor) or mABD-E4S (mABD invalidated for ankG interaction, see Figure 6B). 6 days later (14 div), neurons were fixed and immunostained for GFP, map2 and ankG. Arrowheads indicate the AIS of transfected neurons. **I:** Ratio of the mean fluorescence intensity for GFP labeling in the proximal axon compared to the mean of three dendrites (D) in each transfected neuron (mABD expressed from 0 to 7 div from Fig. 6F for comparison, mABD expressed from 8 to 14 div 1.15 ± 0.09, n=66, 2 independent experiments). **J:** Ratio of the mean fluorescence intensity for ankG labeling in the proximal axon compared to the mean of three dendrites of the same transfected neuron (mGFP 10.79 ± 0.73, n=30; mABD 6.64 ± 0.57, n=114; ABD 11.16 ± 0.51, n=58; mABD-E4S 16.83 ± 0.95, n=66; 2−4 independent experiments).

To functionally confirm that the Nav depletion occurred in these neurons having a morphologically normal AIS, we performed electrophysiological experiments on neurons co-expressing shNav and mABD (Figure 7D & 7E). In these neurons, no INaT could be recorded (7/7 neurons). More remarkably, INaP was either absent (4/7 neurons) or extremely small with an average amplitude as small as 42, 3±21, 5 pA (3/7 neurons) (Figure 7D & 7E). In the 7 slices from which these mABD positive neurons were patched, we also recorded Na+ currents from non-transfected neurons in which the electrophysiological parameters of both INaT and INaP were similar to the currents recorded in control neurons as shown in Figure 2. These results demonstrate that a morphologically normal AIS can be maintained in neurons devoid of Na+ currents.

To confirm the ectopic localization of ankG observed in organotypic slices, we assessed the effect of overexpressing mABD in cultured mature hippocampal neurons (Figure 7F-J). In these neurons, mABD was also localized in the whole neuron with no preferential accumulation at the AIS (AIS/ dendrite ratio for mABD 1.15±0.09, Figure 7F & 7I). mABD over-expression also produced a mislocalization of ankG, that appeared in the cell body and proximal dendrites in addition to the AIS (AIS/dendrite ratio from 10.79±0.73 in control neurons to 6.64±0.57 in mABD-expressing neurons, Figure 7F & 7J). We determined if the two properties of the mABD construct (membrane association and ankG binding) were necessary for this delocalization of ankG. The ABD construct devoid of the farnesylation motif was expressed in a non-polarized manner, and was not able to mislocalize ankG (Figure 7G & 7J). Similarly, the mABD-E4S mutant that lacks ankG binding was localized in a non-polarized manner, and did not delocalize ankG from the AIS (Figure 7H & 7J). Overall, the expression of mABD in mature neurons from slices and cultures shows that binding of ankG to its membrane partners is important for its stabilization at the AIS. Furthermore, the ectopic localization of ankG after mABD expression suggests that ankG binding by mABD is sufficiently strong to perturb ankG targeting, and that the association of ankG with its membrane partners occurs upstream of their insertion into the AIS scaffold.

## Discussion

The AIS is a highly specialized neuronal compartment that plays a key role in neuronal development and excitability. Many studies have highlighted the central role of ankG in the establishment and maintenance of the AIS molecular scaffold, since it targets and anchors most, if not all, known AIS components. However, the converse role of the AIS components on ankG trafficking and stabilization is still poorly understood. Here, we have demonstrated that Nfasc186 and Nav are required for ankG anchoring at the AIS in both developing and mature neurons. Thus, although ankG is unquestionably necessary for AIS construction and stabilization, it is not sufficient by itself to ensure the complete assembly of the AIS.

Both Nav and Nfasc186 are membrane proteins that bind directly to the MBD of ankG via their ankyrin binding domains. Their individual knockdowns produced similar inhibitions of ankG concentration at the AIS, whereas their simultaneous depletions had a cumulative effect (although their ankyrin binding domains are different). This indicates that the ability of Nav and Nfasc186 to anchor ankG to the plasma membrane is crucial for ankG targeting and maintenance at the AIS. Furthermore, the ankG downregulation induced by Nfasc186 depletion is rescued by expressing recombinant Nav channels. This demonstrates that, regardless the identity of the anchoring membrane partner, membrane association is a crucial step in ankG targeting. Such a role of the association between ankG and Nav or Nfasc186 is consistent with recent data from Wang and collaborators who resolved the MBD structure. Indeed, mutations in the MBD, which impair binding to Nav or Nfasc186, also prevent the correct targeting of recombinant ankG to the AIS of cultured hippocampal neurons (Wang et al, 2014). We have not tested whether other AIS membrane partners contribute to ankG targeting. Nevertheless, we can suspect that KCNQ2/3 (Kv7.2/ Kv7.3) channels, which have an ankyrin binding domain very similar to that of Nav channels, could have a similar stabilizing effect (Pan et al, 2006; Hill et al, 2008; Xu & Cooper, 2015).

To pinpoint the features required for ankG stabilization by its membrane partners, we designed a minimal chimeric protein consisting in a membrane-anchored ankyrin binding domain (mABD). This chimeric protein had a dominant negative effect on the concentration of endogenous Nav channels at the AIS. This effect was confirmed by our electrophysiological data obtained in cultured organotypic cortex slices. In this model, neurons expressing shNav alone exhibited a residual INaP (likely produced by Nav1.6 that is not targeted by our shRNA) that was suppressed in neurons co-expressing shNav and mABD. Although mABD expression had a dominant negative effect on endogenous Nav, it rescued the ankG downregulation induced by the shNav and drove the reestablishment of an AIS morphologically indistinguishable from the AIS of naive neurons (see Figure 7B). This demonstrates that properly assembled AIS can be reconstituted by mABD in neurons devoid of Na+ currents. Thus, the membrane-anchoring effect of the Nav channels is essential for ankG targeting to the AIS independently of their ability to produce Na+ current.

Our study unravels two mechanisms regarding AIS components assembly and maintenance. First, we evidenced an unexpected high degree of plasticity of the AIS structure in young neurons. Indeed, when mABD is overexpressed during AIS formation, a larger amount of ankG is positioned to the proximal axon, promoting a wider AIS. These observations suggest that the diameter of the AIS is subject to a plasticity modulated by the amount of available membrane components. These data fit with the observation that the level of Nav channels expressed to the membrane is regulated by a combination of mechanisms involving numerous Nav interacting partners such as ankG, CK2, VGSC beta subunits, FGF13- and 14 (Montersino et al, 2014; Hien et al, 2014; O’Malley & Isom, 2015; Pablo et al, 2016). This AIS diameter plasticity might profoundly influence electrogenesis properties, as shown for AIS length and position plasticity in developing neurons (Galiano et al, 2012; Kuba et al, 2014; Gutzmann et al, 2014). On the opposite, in mature neurons the assembled AIS exhibit very little plasticity unless they are destabilized. Indeed, overexpression of mABD in mature cortical neurons leads to an ectopic localization of ankG that is rescued by a concomitant knockdown for Nav channels. Our data are consistent with the observed stability of AIS components in mature neurons (Hedstrom et al, 2008; Akin et al, 2015) and the preferential regulation of channels immobilization by CK2 in young neurons (Brachet et al, 2010).

A second important finding is the fact that ankG can interact with some of its partners outside the AIS. Indeed, the ability of mABD to mistarget ankG in mature neurons suggests that ankG and its membrane partners, in particular Nav channels, can physically interact with each other upstream of their insertion into the AIS scaffold, and are likely co-transported to the AIS. The mechanisms for the targeting of AIS proteins are the subject of an ongoing debate. On the one hand, a “diffusion trapping” model has been proposed in which Nav and KCNQ2/3 channels are transported to the plasma membrane, diffuse to the AIS where they are immobilized by ankG (Leterrier et al, 2011a; Xu & Cooper, 2015). On the other hand, a “direct insertion” model was put forward, where Nav channels are directly targeted to the AIS where they are immediately immobilized (Akin et al, 2015; Barry et al, 2014). Our model of a co-transport of ankG associated to its membrane partners is in line with the latter hypothesis. We propose that ankG is targeted to vesicular membranes by a tight association with its membranous partners. In our model, the ankG/membrane partner complexes are co-transported to the axon, presumably via kinesin-1 (Barry et al, 2014), and inserted into the AIS scaffold during AIS formation and maintenance.

## Materials and Methods

### Antibodies and plasmids

A rabbit anti-ßIV spectrin antibody (Eurogentec) was developed against amino acid residues 15−38 of the human sequence (XP 006723369). Mouse monoclonal antibodies to ankyrin G (1:400, N106/36 NeuroMab), Nfasc186 (1:200, L11A/41 NeuroMab) and sodium channels (pan Nav; 1:100; Sigma-Aldrich), rabbit polyclonal antibodies to GFP (1:1000; A11122 ThermoFisher), and chicken polyclonal antibodies to map2 (1:10,000; Abcam) were used. Secondary goat antibodies conjugated to Alexa Fluor 488, 555, 647 (ThermoFisher) or DyLight 405 (Jackson ImmunoResearch Laboratories) were used at 1:400 dilutions.

The nucleotide sequences of rat Nav1.2 fragment coding from amino acid 1081 to 1203 (WT, mutated for E1111, mutated for S1112/1123/1124/1126 or mutated for both E and the four S) were obtained by PCR amplification (Expand High Fidelity Taq polymerase, Roche Molecular Biochemicals) on pKv2.1-Nav1.2 plasmids (Brechet et al, 2008) and inserted into the pEGFP-F (Clontech) in the 5’ of EGFP sequence. The resulting sequence encodes a chimeric protein we called mABD (for membranous Ankyrin Binding Domain)(see Figure 6A & 6B). All constructs were verified by DNA sequencing (Beckman Coulter Genomics).

### Animals, cultured hippocampal neurons and organotypic slice culture

All procedures were in agreement with the European Communities Council directive (86/609/EEC). Experiments were performed on pregnant females Wistar rats (Janvier labs) or on rat pups of 7 days. Pregnant females were sacrificed by decapitation and E18 embryos brains were used for primary neuronal cell culture. Primary hippocampal neurons were prepared according to the Banker-type culture protocol (Kaech & Banker, 2006) and transfected at 8 div using Lipofectamine 2000 (ThermoFisher), or nucleofected before plating using an Amaxa rat nucleofector kit (Lonza) according to the manufacturer’s intrusions. P7 rat cortical slice (350 μm) were cultured for 7 days according to the protocol described by Stoppini et al, (1991). At the first day in culture slice were micro-injected with 0.1 to 0.2 μl of lentivirus (titer from 108 to 1010 pfu/ml).

### Lentivus vectors

The pFUGW plasmid (Lois et al, 2002) was modified to express farnesylated EGFP (from pEGFP-F, Clontech), tdTomato (from ptdTomato-N1, Clontech) or mABD (see above) rather than EGFP. On the 5’ of the ubiquitin-C promoter (PacI site) specific shRNA expression cassette (U6/promoter/T5/shRNA/T5/H1promoter from pFIV-H1/U6 siRNA vector, System Biosciences) was introduced by restriction. The sequence of shRNAs directed against ankG, Nav1 and Nfasc186 have been previously described (Hedstrom et al, 2007). Luciferase shRNA (shLuc 159 also know as SHC007 sigma) was validated as a control (Abad et al, 2010). Modified plasmids were either used directly by transfection of neuronal cells or used to produce pseudotyped lentivirus according to (Dull et al, 1998). Lentivirus liberated in cell culture medium was concentrated by ultracentrifugation, titrated and kept at −80° in PBS-1% glycerol. The lentivirus production was performed in the lentivectors production facility/ SFR BioSciences Gerland - Lyon Sud (UMS3444/US8).

### Immunocyto- and immunohistochemistry

Transfected cells were processed for immunofluorescence 6 or 7 days post transfection. Cells were fixed 10 min in 4% paraformaldehyde and then blocked and permeabilized with 0.1% of Triton X-100 and 0.22% gelatin in 0.1 M phosphate buffer for 30 min. Cells were incubated for 1 h with primary antibodies in blocking solution. Corresponding secondary antibodies conjugated to Alexa Fluor or DyLight fluorophores were incubated for 1 h. Coverslips were mounted in Fluor Save reagent (EMD). Free-floating organotypic slices kept on hydrophilized PTFE membranes (Millipore) were fixed for 30 min in 4% paraformaldehyde 7 days post infection. Slices were blocked and permeabilized overnight in PBS containing 0.5% of Triton X-100, 1% Normal Goat Serum (NGS) and 100 μg/ml of Bovine Serum Albumin (BSA). Primary antibodies in PBS containing 0.25% of Triton X-100, 0,5% NGS and 50 μg/ml of BSA were incubated for 4 h. Corresponding secondary antibodies conjugated to Alexa Fluor fluorophores were incubated for 1 h. Slices were counterstained for nuclei with Hoechst 33342 (1 μg/mL) and dry-mounted in ProLong Gold (Life Technologies).

### Images acquisition and analysis

For organotypic slices, image acquisition was performed on a Zeiss LSM780 (Zeiss, Jena, Germany) confocal microscope equipped with a 63X 1.4 N.A. oil immersion objective. Three dimensional z-stacks were collected automatically as frame by frame sequential image series (80 to 120 optical slices). For cultured hippocampal cells, image acquisition was performed on a Zeiss Axio Imager Z2 equipped with a 40X 1.4 N.A. oil immersion objective. For illustration purposes, image editing was performed using ImageJ software (http://rsb.info.nih.gov/ij/) and was limited to Sigma Plus Filter, linear contrast enhancement and gamma adjustment. Quantification was performed using ImageJ. Regions of interest corresponding to AIS or dendrite were manually selected on ankG or GFP images and reported on other channels for intensity measurements. All intensities were corrected for background signal. Fluorescence intensity ratio was calculated for each image individually to circumvent variations related to the immunolabeling. Results are expressed as mean ± SEM. AIS/dendrite ratio is the mean labeling intensity of proximal axon divided by the mean labeling intensity of three dendrites. Polarity index (for ankG quantification in organotypic slice) is the ratio of mean labeling intensities: (AIS - soma) divided by (AIS + soma). The determination of AIS length and width was performed on the ankG labeling using a custom ImageJ script (Leterrier et al, 2015). The beginning and the end of the AIS were determined at 33% of the maximum intensity (Grubb & Burrone, 2010). The width was measured perpendicularly to the axon at the point of maximum intensity of ankG along the AIS. Statistical analyses were performed using Prism 5 (Graphpad Software, La Jolla, CA, USA). Significances were tested using two-tailed unpaired t tests (two conditions) or one-way ANOVA followed by Kruskal-Wallis post-test (three or more conditions). In all figures significance is coded as: *p < 0.05; **p < 0.01; ***p < 0.001.

### Electrophysiology and data analysis

7 or 8 days post transduction organotypic cortical slices were transferred into a recording/perfusion chamber placed on the fixed-stage of an upright microscope (BX51WI, Olympus; fitted with x4 air and x40 water-immersion objectives, a Photonics VX55 camera and a BX-FLA illumination system) and superfused at 1−3 ml/ min with artificial cerebrospinal fluid (ACSF) maintained at room temperature and saturated with 95% O2 and 5% CO2. Voltage-clamp recordings were performed with patch pipettes pulled from borosilicate glass (World Precision Instruments) and having an electrode resistance of 1,5−2,5 MΩ. Na+ currents were isolated by using blockers of K+ and hyperpolarization-activated cation currents (TEA, 4-AP, and Cs) and by omitting Ca2+ from the extracellular solution. Selective recording of Na+ currents was performed by using patch pipettes filled with an internal solution containing (in mM) 140 CsFl (to block K+ and hyperpolarization-activated cation currents), 1 MgCl2, 10 Hepes, 2 EGTA, 2 ATP, 0,2 GTP (pH7,4) and a modified ACSF solution (Na+ current isolation solution) containing (in mM): 120 NaCl, 3 KCl, 26 NaHCO3, 10 Glucose, 1 MgCl2. This external solution did not contain CaCl2 (to prevent Ca2+ currents) but was added with 10 mM tetraethyl-ammonium, 4 mM 4-Aminopyridine, 3 mM kynurenic acid (Sigma Aldrich); to attenuate K+ currents and glutamate synaptic transmission). Separate recording sessions were also performed with 500 nM TTX (Tetrodotoxin, Ascent Scientific) added to the Na+ current isolation solution in order to check that the recorded current was carried by Na+ ions. Na+ currents were evoked by voltage ramps reaching +40 mV from a holding potential of −90 mV and applied at a speed of 0,2 mV/ms.

All recordings were performed exclusively from GFP positive neurons using an Axopatch 200B (Molecular Devices). A computer interfaced to a 12-bit A/D converter (Digidata 1322A using Clampex 9.x; Molecular Devices LLC) controlled the voltage-clamp protocols and data acquisition. The signals were filtered at 5 KHz and digitized at 20 KHz. Uncompensated series resistance was 8±0.5 MΩ. Analysis was conducted in Clampfit 10.3 (Molecular Devices) and SigmaPlot 12 (Systat Software Inc.). After off line linear leak subtraction, the following parameters were measured: threshold voltage for both INaP and INaT, peak voltage for INaP and peak amplitude for both INaP and INaT. Membrane potentials are uncorrected for the liquid junction potential (8,9 mV). Data are shown as mean ± SEM. Mann-Whitney rank sum test was used to test for statistical differences with a significance level of P<0.05.

## Acknowledgements

We thank Dr. Sulayman Dib-Hajj for providing the Nav1.6-GFP plasmid; Aziz Moqrich for critical reading of the manuscript; Hélène Babski and Constance Manso for technical help; Gisèle Froment, Didier Nègre and Caroline Costa (lentivectors production facility /SFR BioSciences Gerland - Lyon Sud (UMS3444/US8) for the production of lentivirus. This work was supported by the Centre National pour la Recherche Scientifique, by a grant to B.D. from the French Agence Nationale de la Recherche (ANR-2011-BSV4-001-1) and by a grant from Conseil Regional PACA (N°2011-10925)

## Author contributions

CL, NC and FC: Conception and design; Experiments and data acquisition; Analysis and interpretation of data; Draft and revision of the article. FR: Experiments and data acquisition AM: Experiments and data acquisition; Analysis and interpretation of data. BD: Conception and design; Draft and revision of the article.

